# Visual activity enhances neuronal excitability in thalamic relay neurons

**DOI:** 10.1101/2023.06.06.543854

**Authors:** Maël Duménieu, Laure Fronzaroli-Molinieres, Cécile Iborra-Bonnaure, Anushka Wakade, Emilie Zanin, Aurore Aziz, Norbert Ankri, Salvatore Incontro, Danièle Denis, Romain Brette, Béatrice Marquèze-Pouey, Dominique Debanne, Michael Russier

## Abstract

The dorsal lateral geniculate nucleus (dLGN) has long been held to act as a basic relay for visual information travelling from the retina to cortical areas, but recent findings suggest a largely underestimated functional plasticity of dLGN neurons. However, the cellular mechanisms supporting this functional plasticity have not been yet explored. In particular, it remains to elucidate whether intrinsic neuronal excitability change upon visual stimuli. We show here that monocular deprivation for 10 days reduces the intrinsic excitability of dorsal LGN relay cells. Furthermore, dLGN neurons exhibit long-term potentiation of their intrinsic excitability (LTP-IE) when suprathreshold afferent retinal inputs are stimulated at 40 Hz or when spikes are induced with direct somatic current injection to reproduce patterns of retinal activity. LTP-IE is observed after eye opening and requires calcium influx mediated by L-type calcium channels. It involves activation of PKA and is expressed through the down-regulation of Kv1 potassium channels. In conclusion, our study provides the first evidence for intrinsic plasticity in dLGN relay cells, thus further pointing the role of thalamic neurons in activity-dependent visual plasticity and amblyopia.

## Introduction

Activity-dependent plasticity in the visual system is classically thought to be exclusively expressed at the cortical level whereas the dorsal lateral geniculate nucleus (dLGN), a primary recipient structure of retinal inputs at the thalamic level, is traditionally considered to be just a relay of visual information. In fact, monocular deprivation has been thought for a long time to produce no effect on the receptive field properties of dLGN neurons (Wiesel and Hubel, 1963). However, this pioneering conclusion has been challenged by later works indicating that this simplistic view does not hold longer (Sherman, 2007; Rose and Bonhoeffer, 2018; Duménieu et al., 2021). The spatial resolution of dLGN neurons activated by the deviate eye in kittens reared with a squint is considerably reduced compared to that of neurons activated by the normal eye (Ikeda and Wright, 1976). In amblyopic patients, functional deficits in visual response are already observed at the stage of the LGN (Hess et al., 2009). Moreover, in contrast to what was previously established, about half of dLGN relay neurons in a given monocular territory receive inputs from each eye, indicating a potential binocularity for large proportion of dLGN neurons (Hammer et al., 2015; Morgan et al., 2016; Rompani et al., 2017). In addition, monocular deprivation was shown to produce a large shift in ocular dominance in dLGN neurons (Jaepel et al., 2017; Sommeijer et al., 2017; Rose and Bonhoeffer, 2018). However, the cellular mechanisms underlying functional changes occurring in these *in vivo* studies remain unclear. In particular, the precise locus of the plasticity has not been clearly identified.

Among the cellular mechanisms that may occur during functional plasticity, regulation of intrinsic neuronal excitability is a potential candidate. Indeed, many brain regions including visual areas express intrinsic plasticity (Mozzachiodi and Byrne, 2010; Debanne et al., 2019). Long-term intrinsic plasticity has been reported in central neurons following stimulation of afferent glutamate inputs (Sourdet et al., 2003), spiking activity (Cudmore and Turrigiano, 2004), sensory stimulation (Aizenman et al., 2003) or following sensory deprivation (Nataraj et al., 2010; Brown et al., 2019). But it is not yet known whether dLGN relay cells also express intrinsic plasticity.

We show here that monocular deprivation reduces intrinsic excitability of dLGN neurons. In addition, we report the induction of long-term potentiation of intrinsic excitability (LTP-IE) in dLGN relay neurons following stimulation of the retinal afferent inputs that evoke spiking activity. This enhanced excitability is observed in a specific developmental window (i.e., after eye opening). LTP-IE is mediated by the down-regulation of Kv1 channels through the activation of PKA. Our results thus provide unambiguous demonstration of the existence of activity-dependent plasticity of intrinsic neuronal excitability in dLGN relay neurons.

## Results

### Monocular deprivation reduces intrinsic excitability in dLGN neurons

Longs Evans rats were monocularly deprived just before eye opening (i.e., at P12) during 10 days and acute slices containing the dLGN were obtained (**Figure 1A**). Contralateral projection zones were identified by injection of cholera toxin conjugated with Alexa 594 and Alexa 488 in separate experiments (**Figure S1A**). Relay cells were recorded in whole-cell configuration from the monocular segment of the contralateral projection zone to the closed eye or from the monocular segment of the contralateral projection zone to the open eye of the dLGN. Compared to neurons activated by the open eye, neurons activated by the deprived eye were less excitable and exhibited a higher rheobase when the neuron was held at a membrane potential of −56 mV to inactivate the T-type calcium current (**Figure S1B**) (deprived side: 82 ± 6 pA n = 13 cells in 10 rats vs. open side: 51 ± 7 pA, n = 10 cells in 9 rats; Mann-Whitney test, p<0.05; **Figure 1B & 1C**; **Figure 1SD**). No significant change in the spike threshold (deprived side: −40.1 ± 1.0, n = 13 vs. −38 ± 0.8 mV, n = 10 on the open side, Mann-Whitney test p>0.1; **Figure S1C**), holding current (deprived side: 30 ± 6 pA, n = 13 vs. 18 ± 6 pA, n = 10 on the open side; Mann-Whitney p>0.5) nor in input resistance (deprived side: 441 ± 69 MΩ, n = 13, open side: 548 ± 46 MΩ, n = 10, Mann-Whitney test, p>0.1; **Figure S1C**) was observed between deprived and spared neurons. Nevertheless, the temporal pattern of the spike discharge was found to be different in deprived and spared neurons. While deprived neurons displayed a reduction in the second inter-spike interval (ISI 2) compared to the first (i.e., ISI 1), neurons activated by the open eye did not show this reduction (deprived side: −83.4 ± 19.8 ms, n = 13 vs. open side: 13.5 ± 18.1, n = 10; Mann-Whitney, p < 0.05; **Figure 1D**). This difference was due to the ramp measured on subthreshold traces (**Figure S1E**). Our results indicate that visual activity promotes neuronal excitability while visual deprivation reduces intrinsic excitability in dLGN neurons.

**Figure 1.**
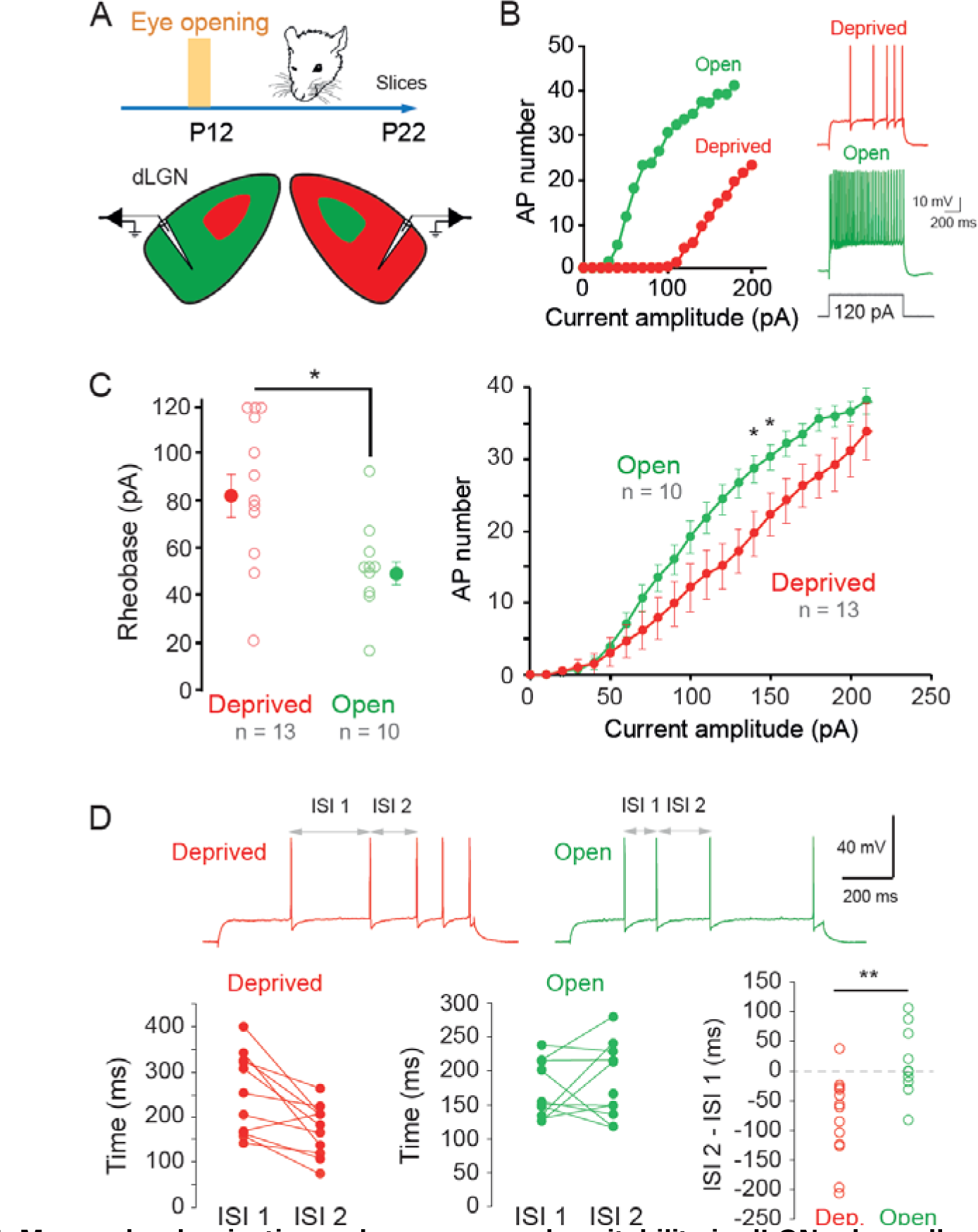
Monocular deprivation reduces neuronal excitability in dLGN relay cells. A. Scheme of the experimental protocol. Lids on one eye were sutured in Long Evans rats at P12 (i.e., just before eye opening). Animals were kept until P22 and acute slices containing both dLGN were prepared. B. Left, comparison of input-output curves from neurons recorded in the contralateral segments of the dLGN activated by the open (green) and deprived (red) eye. Note the difference in rheobase and in intrinsic excitability. Right, comparison of the firing in response to the same current in the two cells. C, Left, comparison of the rheobase in deprived and open dLGN neurons. Right, pooled input-output curves for deprived and open dLGN neurons recorded at −56 mV. *, Mann-Whitney test, p<0.05. D, top, traces from neurons activated by the deprived (red) and open (green) eye to study the adaptation of the first 3 spikes. Bottom left, in neurons activated by the deprived eye, a clear reduction of ISI 2 compared to ISI 1 is observed. Middle, in neurons activated by open eye, no change is observed. Right, comparison of the difference in ISI for neurons activated by the deprived and open eye. **, Mann-Whitney test, p<0.01.

### Induction of long-term potentiation of intrinsic excitability (LTP-IE) in dLGN relay cells

We next checked whether activation of retinal inputs could induce plasticity of intrinsic neuronal excitability during the recording of a dLGN relay neuron. Whole-cell recordings from dLGN relay cells were obtained in acute slices from young rats (P19-P25) at a membrane potential of −65 mV and a stimulating electrode was placed on the optic tract to evoke an excitatory-post-synaptic potential (EPSP; **Figure 2A**). GABA_A_ receptors were blocked with picrotoxin (100 µm). EPSP amplitude increased with stimulus intensity and evoked an action potential (**Figure S2A**). Long-term potentiation of intrinsic excitability (LTP-IE) measured 20-30 minutes after the stimulation was induced in dLGN relay cells following stimulating supra-threshold EPSP (i.e., producing an action potential on the top of each EPSP) at a frequency of 40 Hz during 10 minutes. The number of spikes evoked by a same pulse of current increased by a factor two (196 ± 19%, n = 11; mean spike number: 7.1 ± 0.8 before and 13.3 ± 1.2 after stimulation, Wilcoxon test p<0.001; **Figure 2B**). This plasticity was not associated with any change in input resistance (103 ± 1%, n = 11; mean resistance: 369 ± 28 MΩ before and 373 ± 21 MΩ after stimulation, Wilcoxon test p>0.5; **Figure 2B**, **Figure S2C**), nor in holding current to maintain the membrane potential at −65 mV (**Figure S2B**). The presence of spikes during the induction was found to be critical in the magnitude of LTP-IE. When subthreshold EPSPs were evoked, the magnitude of LTP-IE was significantly smaller (130 ± 6 %, n = 8) than that produced by suprathreshold EPSPs eliciting action potentials (196 ± 19%, n = 11, Wilcoxon test p<0.05; **Figure 2C**). Here again, input resistance was unchanged after stimulation of subthreshold EPSPs (**Figure S2D**). Furthermore, subthreshold EPSPs did not induce a larger LTP-IE than that observed when the stimulation was suspended during 10 minutes (129 ± 6%, n = 7, Mann-Whitney, p>0.05; **Figure 2C**), indicating that spiking activity is critical for inducing LTP-IE.

**Figure 2.**
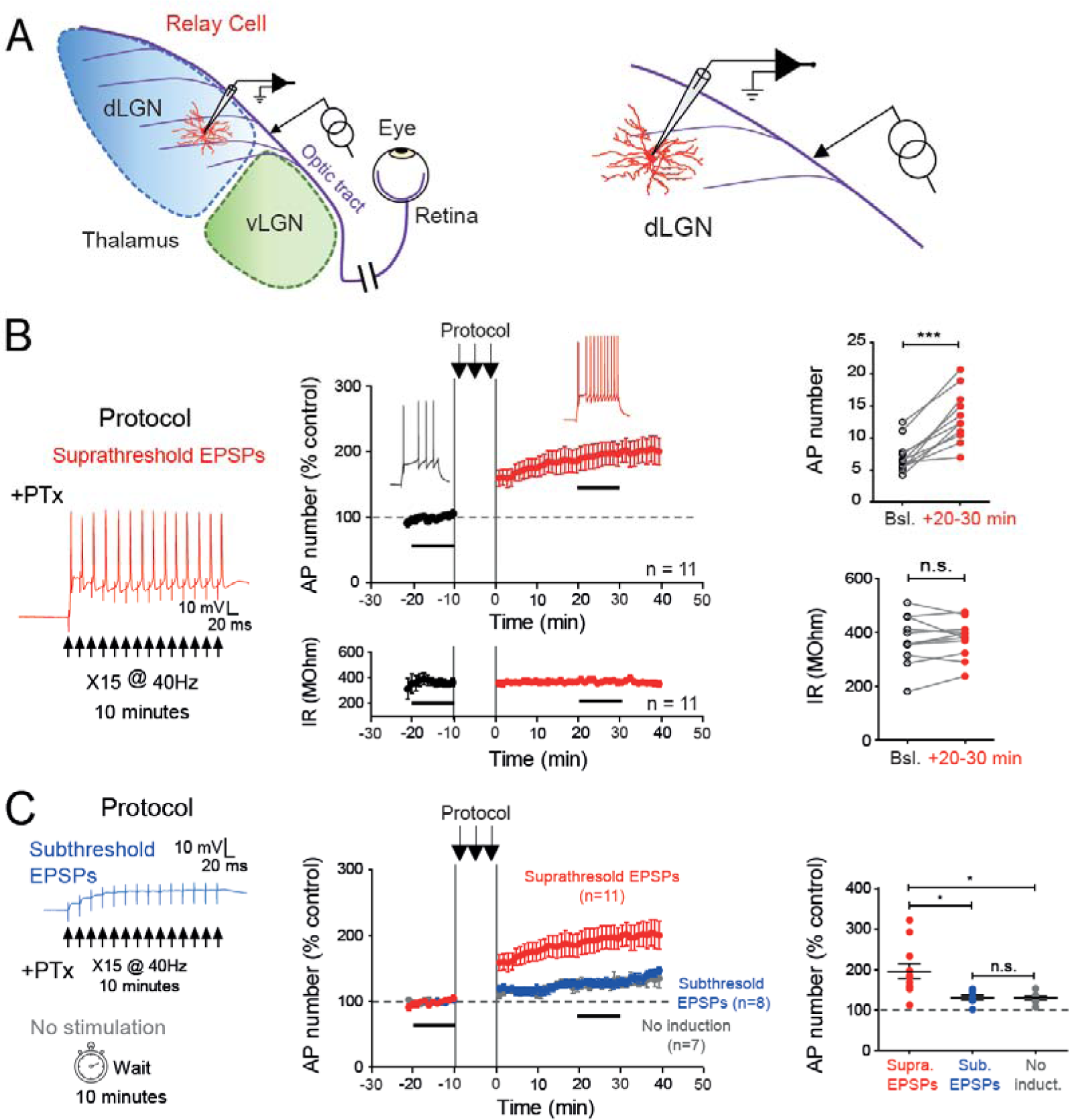
Synaptic induction of LTP-IE in dLGN relay neurons. A. Recording configuration of relay cells in the dLGN. Left, scheme of the LGN with the dorsal LGN (dLGN, in blue), the ventral LGN (vLGN, green) and the afferent axons from the retina (optic tract, violet). Right, recording and stimulation arrangements. B. Trains of suprathreshold EPSPs delivered at a frequency of 40 Hz induce LTP-IE in dLGN relay neurons. Left, protocol. Middle, time-course of the spike number normalized to the control period (top) and time-course of the input resistance (Rin, bottom). Right, comparison of the AP number (top) and the Rin (bottom). Wilcoxon test, ***, p<0.001; ns, not significant. C. Trains of subthreshold EPSPs do not induce LTP-IE. Left, protocol of stimulation with subthreshold EPSPs (top) and without any stimulation (bottom). Middle, Time-courses of excitability changes following stimulation with subthreshold EPSPs (blue) and no stimulation (grey) compared to the time-course of excitability changes following stimulation with suprathreshold EPSPs. Right, comparison of excitability changes in dLGN neurons following suprathreshold EPSPs (red), subthreshold EPSPs (blue) and no stimulation (grey). Mann-Whitney test, *, p<0.05; ns, not significant.

### Action potential firing is sufficient to induce LTP-IE in dLGN relay neurons

In order to check whether spiking activity alone was the key trigger of LTP-IE in dLGN neurons, action potentials were induced by current pulses delivered at 40 Hz in the presence of ionotropic glutamate and GABA receptors antagonists. A two-fold increase in excitability that could not be distinguished from that induced by synaptic stimulation was observed in these conditions (240 ± 22%, n = 15; mean spike number: 4.6 ± 0.5 before and 10.6 ± 1.2 after stimulation, Wilcoxon test p<0.001; **Figure 3A**). R_in_ remained unchanged (396 ± 24 MΩ before vs. 395 ± 22 MΩ after induction, n = 15, Wilcoxon test p >0.1, data not shown). We conclude that spiking activity is critical for induction of LTP-IE in dLGN neurons.

**Figure 3.**
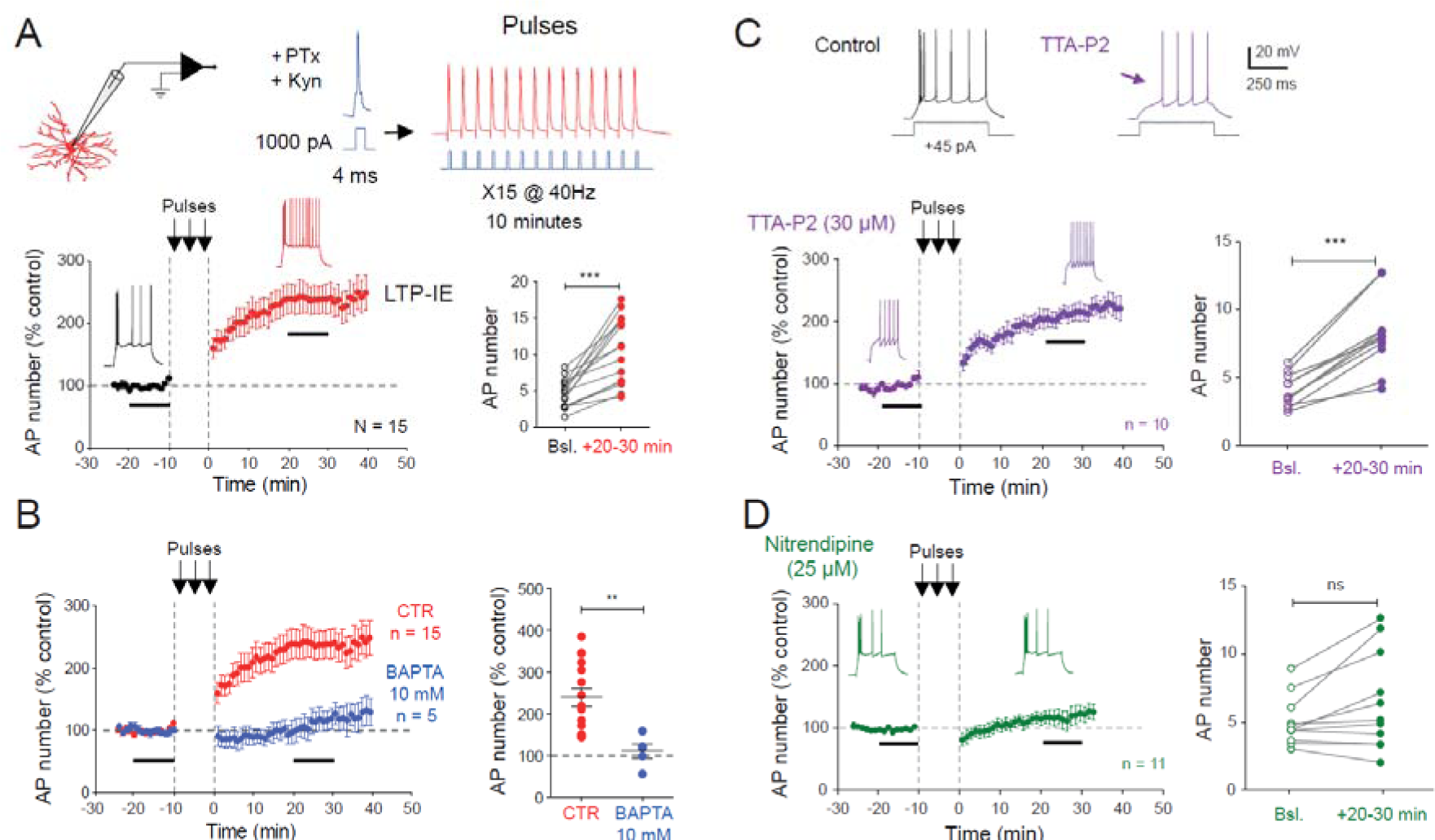
Induction of LTP-IE in dLGN relay neurons with spiking activity. A. Trains of APs at 40 Hz induce LTP-IE in dLGN relay neurons. Top, recording configuration and protocol. Bottom left, time-course of neuronal excitability before and after trains of APs. Bottom right, group data for AP number. Wilcoxon test, ***, p<0.001. Bsl: baseline. Kyn: kynurenate, PTx: picrotoxin. B. Intracellular BAPTA blocks LTP-IE. Left, time-courses of changes in excitability in BAPTA (blue) compared to control (red). Right, comparison of the normalized change in neuronal excitability. Mann-Whitney test, **, p<0.01. C. LTP-IE is not mediated by T-type calcium channels. The T-type calcium channel blocker TTA-P2 suppressed the initial burst and uncovered a ramp-and-delay phenotype. In the presence of TTA-P2, LTP-IE is still induced. Left, time course. Right group data. Wilcoxon test, ***, p<0.001. Bsl: baseline D. Lack of LTP-IE in the presence of the L-type calcium channel blocker, nitrendipine. Left, time course. Right, group data. Wilcoxon test, ns, p>0.1.

We next evaluated whether the triggering signal of LTP-IE relied on an elevation of intracellular calcium. As action potentials activate voltage-gated calcium channels, we tested whether chelating intracellular calcium variations with BAPTA (10 mM) would prevent LTP-IE induction in dLGN neurons. In this condition, no LTP-IE was observed (111 ± 17%, n = 5; **Figure 3B**). Given that dLGN relay cells express a large T-type calcium current (Crunelli et al., 1989; Suzuki and Rogawski, 1989), we first investigated the role of these channels in the induction of LTP-IE. Blockade of T-type channels with the specific Cav3.2 channel blocker, TTA-P2, abolished the depolarizing rebound and the initial burst of action potentials (**Figure 3C**), but left intact the increased neuronal excitability induced by spiking activity at 40 Hz (197 ± 11%, n = 10; mean spike number: 4.2 ± 0.4 before and 8.5 ± 0.9 after stimulation, Wilcoxon test p<0.01; **Figure 3C**). These results indicate that calcium is required for induction of LTP-IE but the source of calcium is not mediated by T-type Ca^2+^ channels. We thus, checked whether L-type (Cav1) Ca^2+^ channels could be involved in LTP-IE induction. In the presence of the L-type Ca^2+^ channel blocker, nitrendipine (25 µM), no LTP-IE was induced by spiking activity (122 ± 11%, n = 11; AP number before: 5.0 ± 1.0 after: 6.4 ± 1.1, Wilcoxon test, p>0.1; **Figure 3D**), indicating that sodium action potentials activate L-type Ca^2+^ channels to trigger the calcium influx leading to LTP-IE.

### LTP-IE can only be induced after eye opening

We next examined the developmental profile of LTP-IE induced by spiking stimulation at 40 Hz in dLGN neurons. For this, we extended the range of age from P19-P25 to P9-P30. Interestingly, while LTP-IE was robustly induced from P17 to P30 (215 ± 12%, n = 39; **Figure 4A**), the magnitude of LTP-IE was much reduced before eye opening (P9-P12; 127 ± 13%, n = 10; **Figure 4A**) and just after eye-opening (P13-P16; 137 ± 7%, n = 12; **Figure 4A**). In fact, in contrast to the situation at P17-P30, the spike number was not significantly increased at P9-P12 (from 3.0 ± 0.3 to 3.9 ± 0.6 spikes, n = 10; **Figure S3A**). These results reveal that LTP-IE is developmentally regulated as it is only robustly expressed after P17, i.e., after eye opening.

**Figure 4.**
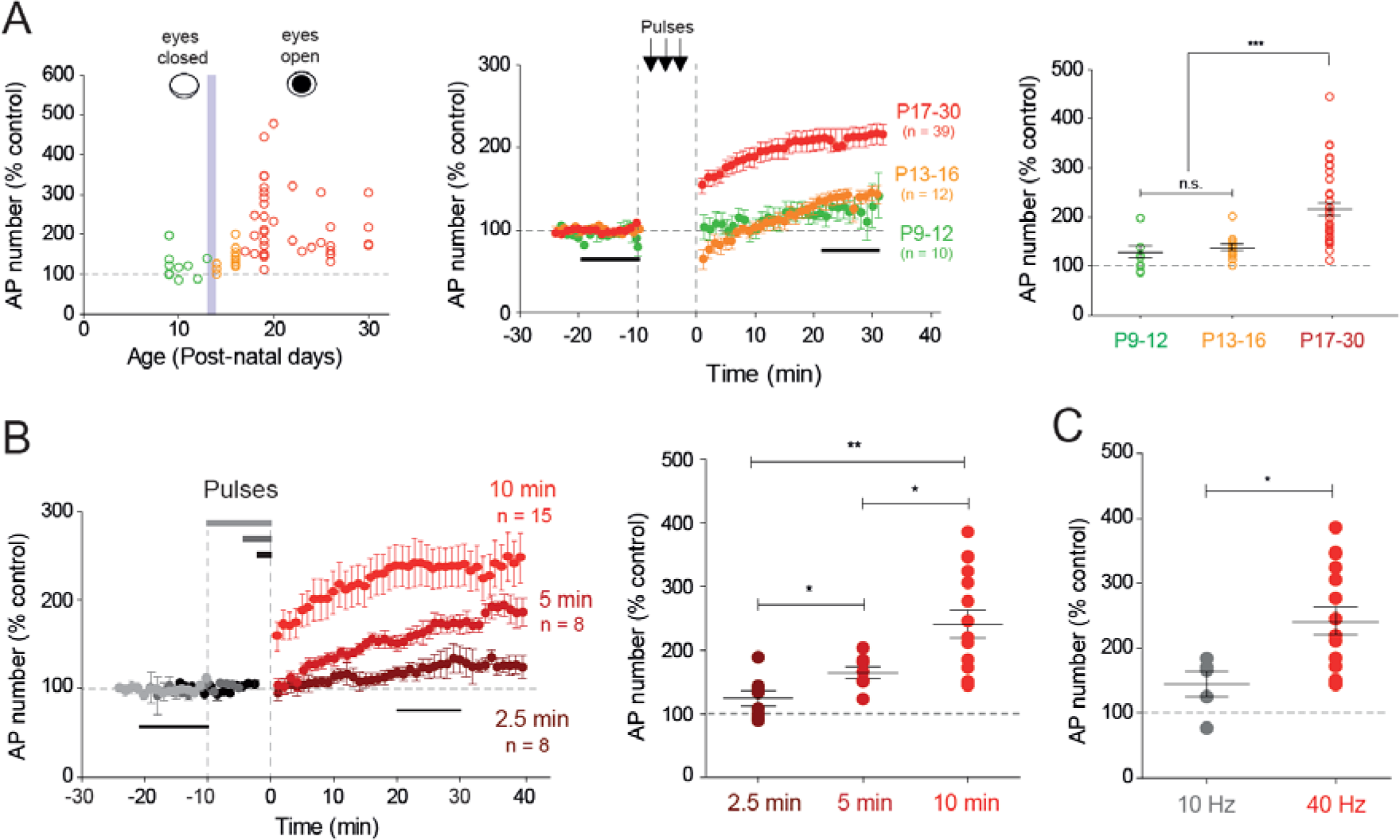
Developmental profile and incidence of stimulus duration and frequency on LTP-IE. A. Left, normalized LTP-IE as a function of the postnatal age. The vertical grey bar represents the time of eye opening. Note that before eye opening, no LTP-IE is induced. Middle, time-courses of LTP-IE induced at P9-P12 (green), P13-P16 (orange) and P17-P30 (red). Right, comparison of LTP-IE at different ages. Mann-Whitney test, ns, not significant; ***, p<0.001. B. Role of the stimulus duration on LTP-IE in dLGN neurons. Left, time-course of LTP-IE induced by a stimulation lasting 2.5 min (brown), 5 min (dark red) and 10 min (red). Right, comparison of the magnitude of LTP-IE. Mann-Whitney test, *, p<0.05, **, p<0.01. C. Frequency-dependence of LTP-IE. Comparison of the magnitude of LTP-IE induced by stimulation at 10 and 40 Hz. Mann-Whitney test, *, p<0.05.

### Properties of LTP-IE induction in dLGN relay neurons

We next checked whether the duration of stimulation was critical for LTP-IE induction with 40 Hz stimulation by current injection. While no significant LTP-IE was induced with a stimulation time of 2.5 min (124 ± 10%, n = 8; Wilcoxon test p>0.1, **Figure 4B**), a significant LTP-IE was induced with a stimulation time of 5 min (164 ± 9%, n = 8; Wilcoxon test p<0.01; **Figure 4B**). Nevertheless, the magnitude of LTP-IE was significantly smaller than those produced by 10 min stimulation (MW, p<0.05; **Figure 4B**).

As dLGN neurons are able to fire at a wide range of frequencies during visual stimulation in rodents, we checked the effect of stimulation frequency on intrinsic excitability. At 10 Hz, no significant facilitation was observed (145 ± 9%, n = 5; **Figure 4C**; mean spike number 5.3 ± 0.7, n = 5 before 10 Hz stimulation, and 7.4 ± 0.9 after 10 Hz stimulation, Wilcoxon test p > 0.05; **Figure S3B**). We conclude that LTP-IE in dLGN neurons is frequency-selective and is preferentially induced by stimulation at 40 Hz.

### Expression of LTP-IE in dLGN neurons involves Kv1 channels

We next determined the expression mechanisms of LTP-IE in dLGN neurons. In the presence of the Cav3.2 channel blocker, TTA-P2, the initial burst disappeared and revealed a ramp-and-delay phenotype that is characteristic of the contribution of Kv1 channels (Cudmore et al., 2010). Furthermore, induction of LTP-IE was associated with a reduction in the first spike latency (50 ± 1%, n = 9; mean latency: 187 ± 4 ms before and 96 ± 3 ms after stimulation, Wilcoxon test p<0.01; **Figure 5A**). Similar changes in the latency of the first spike after the initial T-type calcium burst were also observed in LTP-IE induced either by supra-threshold synaptic stimulation (50 ± 4%, n = 10; mean latency: 184 ± 18 ms before and 92 ± 12 ms after stimulation, Wilcoxon test p<0.01; **Figure S4A**) or by trains of APs induced by direct current injection (63 ± 4%, n = 15; mean latency: 274 ± 22 ms before and 174 ± 21 ms after stimulation, Wilcoxon test p<0.01; **Figure S4B**). In addition, the jitter to the first spike was found to be reduced following LTP-IE induction in the presence pf TTA-P2 (standard error: 38.6 ± 4.2 ms before induction vs. 14.4 ± 1.1 ms, n = 10, Wilcoxon p<0.01; **Figure S5**) and the spike threshold was hyperpolarized (from −36.1 ± 1.7 mV to −39.4 ± 1.3 mV, n = 10, Wilcoxon test, p<0.01; **Figure 5B**). Hyperpolarization of the spike threshold was also seen following induction of LTP-IE with suprathreshold retinal inputs (from −37.4 ± 0.8 mV to −39.2 ± 0.8 mV, n = 10; Wilcoxon test, p<0.01, **Figure S6A**) or current injection (from - 39.6 ± 0.9 mV to −42.8 ± 1.1 mV, n = 15; Wilcoxon test, p<0.001; **Figure S6B**). Such modulations in first spike latency, jitter and threshold are indicative of Kv1 channel down-regulation (Goldberg et al., 2008; Cudmore et al., 2010; Campanac et al., 2013).To verify this hypothesis, experiments were conducted in the presence of the broad spectrum Kv1 channel blocker, DTx-I or in the presence of the specific Kv1.1 channel blocker, DTx-K. As expected, DTx-I (100 nM) or DTx-K (100 nM) was found to mimic LTP-IE (i.e., increase of AP number by a factor ∼3, AP hyperpolarization by ∼4 mV and latency of the first spike after the burst by half; **Figure 5C** & **Figure S7**), but more interestingly LTP-IE was totally occluded in the presence of DTx-I or DTx-K (94 ± 8%, n = 11; mean spike number: 8.4 ± 1.0 before and 8.0 ± 1.2 after stimulation; Wilcoxon test p>0.1; **Figure 5C**). We conclude that LTP-IE in dLGN relay neurons is mediated by the down-regulation of Kv1 channels.

**Figure 5.**
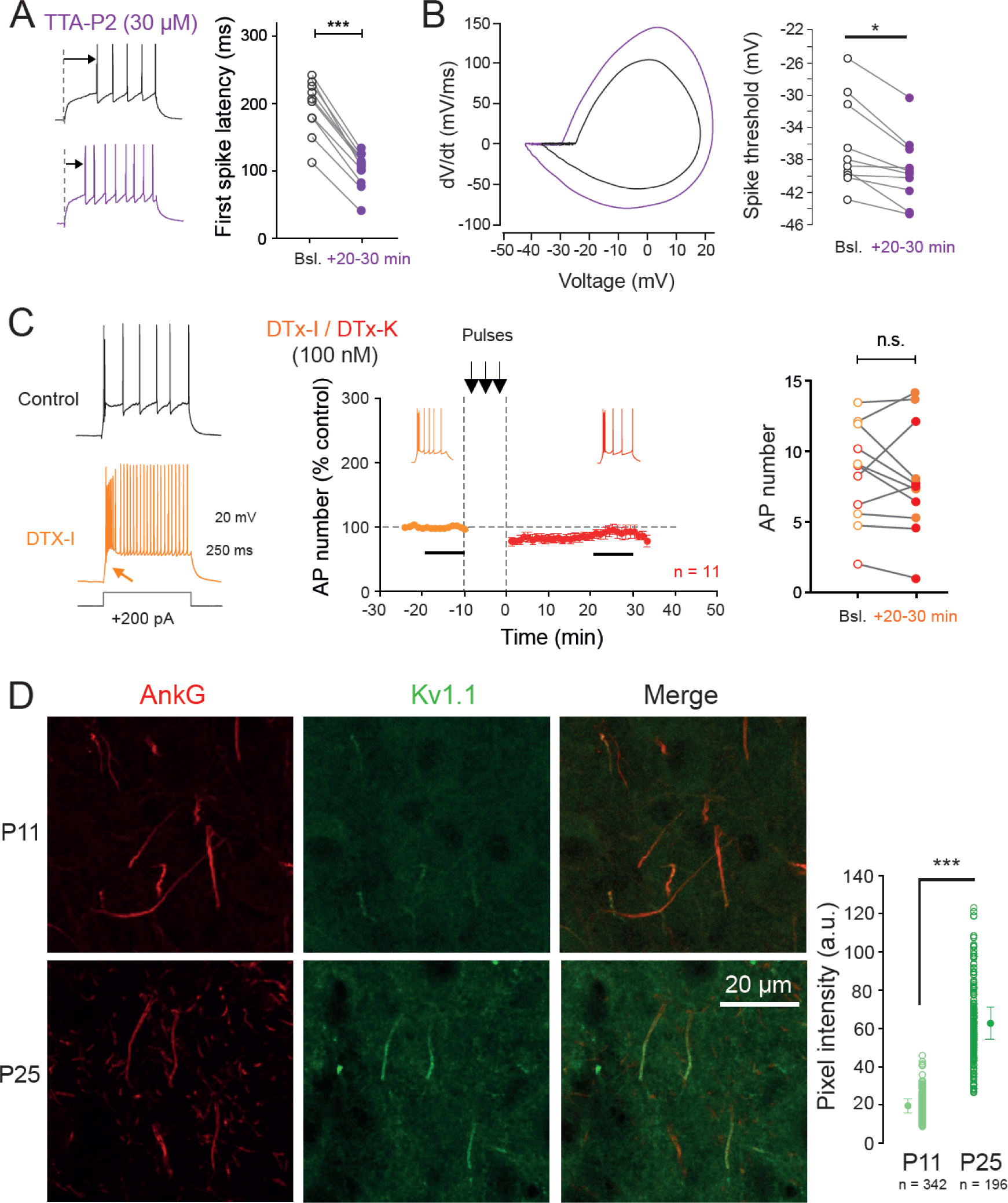
Expression mechanisms of LTP-IE in dLGN relay neurons: Kv1 channels. A. Reduction of the first spike latency in dLGN neurons following induction of LTP-IE in the presence of T-type channel blocker. Left, comparison of the first spike latency before and after induction of LTP-IE in the presence of TTA-P2. Right, group data. Wilcoxon test, ***, p<0.001. B. Hyperpolarisation of the AP threshold. Left, phase plots. Right, pooled data. Wilcoxon test, *, p<0.05. Bsl: baseline. C. Pharmacological blockade of Kv1 channels with DTx-I/K prevents the induction of LTP-IE by 40 Hz spiking activity. Left, effect of DTx-I on a dLGN relay neuron. Middle, time-course of the LTP-IE in the presence of DTx-I/K. Right, group data. Wilcoxon test, ns, not significant. Bsl: baseline. D. Kv1.1 channels are immuno-detected on the axon initial segment of dLGN neurons at P25 (bottom raw) but not at P11 (top raw). Left, column, Ankyrin G labelling. Middle column, Kv1.1 immuno-labelling. Right, column, merge of the Kv1.1 and AnkG immuno-labelling. Right, pooled data. Mann-Whitney test, p<0.001

As LTP-IE induction requires Kv1 channels and LTP-IE is absent before eye opening, we made the hypothesis of a developmental regulation of Kv1 channel expression. For this, we made immunostaining of both ankyrin G, a scaffolding protein of the axon initial segment (AIS) and Kv1.1 channels in thin slices of P11 and P25 rat dLGN. While at P25, Kv1.1 and ankyrin G were co-localized at the AIS, at P11 ankyrin G but not Kv1.1 was stained at the AIS (**Figure 5D**). The quantification of the Kv1.1 immunostaining at the AIS indicate that it increases by a factor 3 between P11 and P25 (**Figure 5D**).

### LTP-IE in dLGN neurons requires PKA activity but not arachidonic acid

Kv1 channels are regulated by many molecules including lipids as arachidonic acid (Oliver et al., 2004; Carta et al., 2014). Inhibition of phospholipase A2 (PLA2; i.e., the arachidonic acid synthesis enzyme) with AACOCF3 (20 µM) did not prevent induction of LTP-IE (177 ± 18%, n = 6; **Figure S8A**), indicating that arachidonic acid is not involved in the down-regulation of Kv1 channels.

In visual cortical neurons, LTP-IE induced by a similar protocol (i.e., spiking activity at 40 Hz during 10 min) requires PKA activity (Cudmore and Turrigiano, 2004). We therefore tested whether LTP-IE in dLGN neurons also involves PKA. Inhibition of PKA with KT5720 (1 µM) prevented LTP-IE (117 ± 10%, n = 8; AP number: 8 ± 1 before vs. 9 ± 2 after, Wilcoxon test, p>0.05; **Figure 6A**), indicating that activation of PKA is required for the induction of LTP-IE. Furthermore, application of the PKA activator forskolin (30 µM) during 10 min in the presence of the HCN channel blocker, ZD-7288 (1 µM) to prevent modulation of h-channels induced a mild LTP-IE (151 ± 11%, n = 8; **Figure 6B**). AP number increased from 6.5 ± 0.7 to 9.6 ± 1.2, n = 8 after forskolin application (Wilcoxon test, p<0.01; **Figure 6B**). Interestingly, here again, a hyperpolarization of the spike threshold (−36.1 ± 0.6 mV vs. −37.1 ± 0.6 mV, n = 8; Wilcoxon test, p<0.05; **Figure S8B**) and a reduction in the first spike latency after the burst (69 ± 6%, n = 8; mean latency: 223 ± 29 ms before and 152 ± 23 ms after stimulation, Wilcoxon test p<0.01; **Figure S8C**) were observed, suggesting that forskolin down-regulates Kv1 channels. Altogether, our results indicate that Ca^2+^ entry by L-type calcium channels activate PKA that in turn reduces Kv1 channel activity (**Figure 6C**).

**Figure 6.**
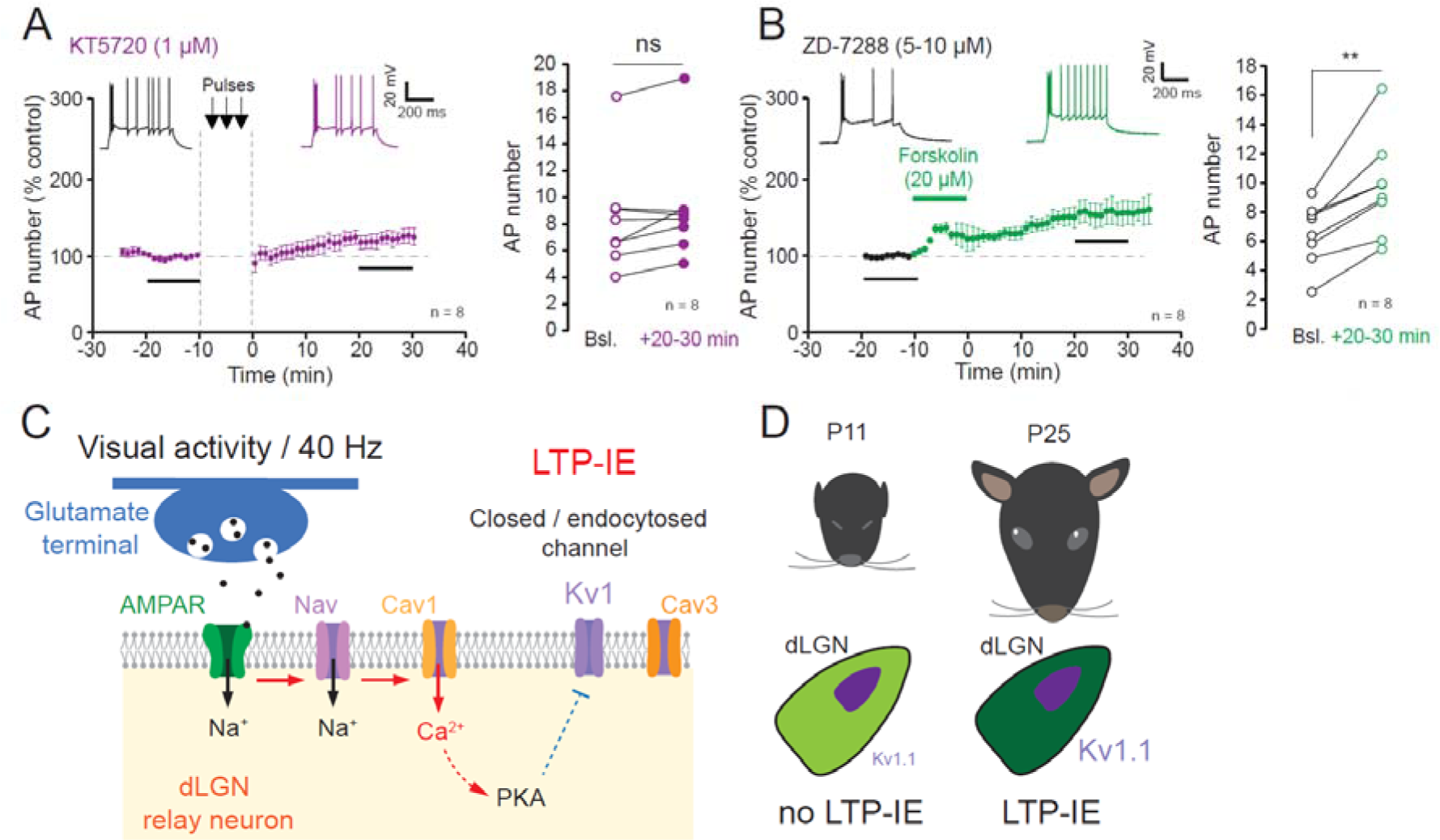
Requirement of PKA and summary. A. Lack of LTP-IE in dLGN neurons recorded in the presence of the PKA inhibitor KT5720. Left, time-course. Right pooled data. Wilcoxon test, ns, not significant. Bsl: baseline. B. The PKA activator, forskolin, induces LTP-IE in dLGN neurons in the presence of the HCN-channel blocker ZD-7288. Left, time course with representative traces. Note the change in the latency of single action potentials after the initial burst. Right, pooled data. Wilcoxon test, ** p<0.01. C & D. Summary of the mechanisms of LTP-IE in dLGN neurons. C. Synaptic release of glutamate from retinal input at 40 Hz activates AMPAR that depolarizes the membrane and activates Nav channels. Sodium action potentials open L-type Cav1 channels provoking a calcium entry that activates PKA. T-type Cav3 channels do not play a role in the induction. Solid red arrows indicate a direct activation whereas the dashed red arrow indicates an indirect activation and the blue connexion represents an indirect inhibition. D. At P11, the eyes are closed and dLGN express little Kv1.1 channels, thus limiting LTP-IE expression. At P25, eyes are open and Kv1.1 are fully expressed allowing induction of LTP-IE.

## Discussion

We show here that monocular deprivation reduces intrinsic excitability in dLGN relay cells of young rats, suggesting for the first time that visual activity enhances neuronal excitability in visual thalamic neurons. In addition, dLGN neurons display long-lasting enhancement of their intrinsic excitability upon stimulation of retinal inputs that elicit supra-threshold spiking activity. LTP-IE in dLGN relay neurons is also induced by spiking activity alone indicating that synaptic activity does not play a critical role in its induction. In addition, LTP-IE depends on a postsynaptic calcium influx, that is not mediated by low-threshold T-type (Cav3) calcium channels but by high threshold L-type (Cav1) calcium channels. Furthermore, we show that LTP-IE in dLGN relay cells involves the down-regulation of Kv1 channels, as first spike latency is reduced and LTP-IE is blocked in the presence of DTx-I or DTx-K. LTP-IE is mediated by PKA activation but not production of AA. LTP-IE could not be expressed before eye opening indicating that it is developmentally regulated. Kv1.1 channel immunostaining was found at the AIS in P25 but not in P11 animals, suggesting that Kv1.1 channels may constitute the limiting factor for the expression of LTP-IE before eye opening. Our findings therefore constitute strong evidence for the presence of activity-dependent plasticity of intrinsic excitability in visual thalamic neurons both *in vitro* and *in vivo*.

### Monocular deprivation reduces intrinsic excitability in dLGN neurons

We show here that monocular deprivation (MD) for 10 days reduces intrinsic neuronal excitability of dLGN neurons receiving the deprived eye inputs compared to dLGN neurons receiving the open eye inputs. A significant change in the rheobase was observed between the two populations of cells but no changes in input resistance nor in AP threshold were observed. This result strongly suggests that visual activity promotes neuronal excitability in dLGN relay neurons according to the Hebbian plasticity scheme. In fact, MD induces a shift in ocular dominance in the dLGN in adult mice (Jaepel et al., 2017).

MD is responsible for a lack of visual acuity on the deprived eye, called the amblyopic eye. This loss of function is classically thought to result from synaptic changes. However, it may also result from changes in intrinsic neuronal excitability. In fact, reduction in excitability following MD has been reported in L5 pyramidal neurons of the visual cortex (Nataraj et al., 2010). Our results therefore support the idea that inactive neurons become less excitable already at the thalamic stage.

### Induction mechanisms of intrinsic plasticity in dLGN relay cells

We show that LTP-IE in dLGN relay neurons was induced by synaptic stimulation at 40 Hz that produced spiking activity but not by subthreshold EPSPs, indicating that synaptic stimulation does not play a critical role in the induction of intrinsic plasticity. In contrast to cortico-thalamic synapses, retino-thalamic synapses are located near the cell body (Sherman, 2016) and generate a large synaptic current able to trigger a postsynaptic action potential (Blitz and Regehr, 2003). The requirement of spiking activity was further demonstrated by the lack of potentiation when no stimulation was delivered and by the fact that spiking activity induced by current steps (i.e., with no synaptic stimulation) induced LTP-IE of similar magnitude.

Spiking activity induces LTP-IE in dLGN neurons through a postsynaptic calcium influx since neurons recorded with a pipette filled with BAPTA displayed no LTP-IE. This calcium influx is not mediated by T-type calcium channels since TTA-P2 did not prevent induction of LTP-IE. Rather, calcium influx is mediated by high-threshold L-type calcium influx activated by action potentials. T-type calcium channels have been shown to occupy a critical position in the induction of synaptic plasticity in subcortical neurons (Pigeat et al., 2015; Leresche and Lambert, 2017). L-type calcium channels are involved in the induction of LTP and LTD in dLGN neurons (Ziburkus et al., 2009) and in AIS plasticity (Evans et al., 2013). In addition, L-type calcium channels are known to play a critical role in retinogeniculate refinement as they contribute to the plateau potential activated by synaptic inputs (Dilger et al., 2015).

### Stimulation parameters: duration and frequency

We show that a minimal duration of 5 minutes is required to induce a significant increase in intrinsic excitability, as no significant enhancement of intrinsic excitability is observed with 2.5 minutes of 40 Hz stimulation. In addition, we show that the lack of stimulation during 10 minutes induces no LTP-IE.

Firing frequency was also found to be critical as a stimulation at 10 Hz did not induce a significant increase in intrinsic excitability in dLGN relay neurons. This may result from the lack of summation of calcium influx at 10 Hz but not at 40 Hz. During visual stimulation, dLGN relay neurons in rodents fire at mean frequencies ranging from ∼5 Hz to ∼50 Hz (Denman and Contreras, 2016; Sriram et al., 2016). Thus, firing frequency at 40 Hz used to induce LTP-IE in dLGN relay neurons corresponds to a physiological firing frequency observed during visual stimulation.

### Expression mechanisms of intrinsic plasticity in dLGN relay cells

Long-lasting intrinsic plasticity in cortical and hippocampal neurons involves a wide set of voltage-gated channels including HCN, SK, Kv1 and Kv7 channels (Sourdet et al., 2003; Campanac et al., 2008, 2013; Debanne et al., 2019; Incontro et al., 2021; Sammari et al., 2022). We show that Kv1 channels are involved in the expression of LTP-IE in dLGN neurons. First, consistent with a loss of function of the Kv1 channels (Campanac et al., 2013), the latency of the first action potential was reduced. Furthermore, the spike jitter was also reduced and the spike threshold was elevated (Cudmore et al., 2010; Campanac et al., 2013). Finally, LTP-IE was occluded by DTx. Down-regulation of Kv1 channel activity does not involve the production of arachidonic acid. In contrast, PKA is involved in LTP-IE induction as its inhibition prevents LTP-IE induction and its activation by forskolin enhances intrinsic excitability. Interestingly, the spike latency was also reduced by forskolin suggesting that PKA also targets Kv1 channels. The loss of function of Kv1 channels by PKA has been shown to be mediated by Kv1 channel endocytosis (Williams et al., 2012).

Whether these expression mechanisms identified in LTP-IE *in vitro* also occur during MD *in vivo* is still unclear. First, LTP-IE is monitored during a few tens of minutes in *in vitro* experiments without any sign of decrement but it is not known whether this persists for hours or days. Second, the activity of thalamic neurons is controlled by many neuromodulators such as serotonin, acetylcholine, noradrenaline, and orexin (Funke et al., 1993; Grubb et al., 2003; Grubb and Thompson, 2004; Orlowska-Feuer et al., 2019; Sokhadze et al., 2022; Reggiani et al., 2023) that may tune neuronal excitability during visual experience.

### Reduction in first spike latency: a possible link between intrinsic and synaptic changes

One of the major functional consequences of LTP-IE in dLGN relay neurons is a reduction in the first spike latency, as a result of a loss of Kv1 function. This effect is visible when the T-type calcium current is inactivated by pharmacological tools (TTA-P2) or by the depolarization caused by various neuromodulators (McCormick and Bal, 1997). The major consequence of the Kv1-dependent latency shortening would be to promote the facilitation of the output synaptic message in the cortex by a mechanism that depends on the minimization of axonal sodium channel inactivation (Zbili et al., 2020). Although thalamo-cortical axons are > 2 mm long in rodent (Little et al., 2009), the presence of myelin prolongs their space constant up to 3 mm, and thus makes possible the modulation of the output message (Zbili and Debanne, 2020). Thus, intrinsic modifications at the stage of the dLGN may have synaptic consequences in the cortex.

### Developmental regulation of LTP-IE

We report here that LTP-IE is developmentally regulated as no LTP-IE is observed before P17. However, it is observed up to P30 with no apparent decline. A similar postnatal onset for plasticity (i.e. P17) has been recently reported for the plasticity of ocular dominance in the mouse dLGN (Li et al., 2023). In addition, a shift in OD has been also reported in adult mouse dLGN (Jaepel et al., 2017). Interestingly, we show that Kv1.1 immuno-labelling at the AIS increases by a factor 3 from P11 to P25, suggesting that expression of Kv1.1 channels at the AIS may represent a limiting factor for the expression of LTP-IE.

### Implication in amblyopia

Amblyopia is characterized by a reduction of visual acuity through the eye that did not function properly during postnatal development. So far, this lack of visual function was thought to be located at the cortical stage. Our findings clearly support the idea that functional changes already occur at the thalamic level as neuronal excitability is significantly elevated in neurons activated by the open eye compared to that of the deprived eye. The subthreshold ramp observed in neurons recorded within the deprived region suggests that firing is delayed for deprived neurons. This delayed firing will promote inactivation of sodium channels that would in turn weaken synaptic outputs at the cortical level (Zbili et al., 2020).

## Methods

### Monocular deprivation & identification of projection zones

All experiments were conducted according to the European and Institutional guidelines (Council Directive 86/609/EEC and French National Research Council and approved by the local health authority (Veterinary services, Préfecture des Bouches-du-Rhône, Marseille; APAFIS n°7661-2016111714137296 v4)). Young (P12) Long Evans rats of both sexes were anesthetized with isoflurane and lids of the right eye were sutured for 10 days after adding eye drops containing a local anaesthetic (tetracaine 1%). Sutures were checked each day and if not intact, animals were not used.

P12 rats were deeply anesthetized with isoflurane and the sclera and cornea were pierced with a syringe containing a 1% solution of cholera toxin B subunit (Invitrogen) conjugated to either Alexa Fluor 488 (green) or Alexa Fluor 555 (red) dissolved in distilled water to inject a different fluorescent marker in each eye (Krahe and Guido, 2011). The animals were kept for 1-2 days after injection.

### Acute slices of rat dLGN

Thalamic slices (350 µm) were obtained from 9- to 30-day-old Long Evans rats of both sexes. Rats were deeply anesthetized with isoflurane and killed by decapitation. Slices were cut in an ice-cold solution containing (in mM): 92 *n*-methyl-D-glutamine, 30 NaHCO_3_, 25 D-glucose, 10 MgCl_2_, 2.5 KCl, 0.5 CaCl_2_, 1.2 NaH_2_PO_4_, 20 Hepes, 5 sodium ascorbate, 2 thiourea, and 3 sodium pyruvate and bubbled with 95% O_2_ - 5% CO_2_ (pH 7.4). Slices recovered (20-30 min) in the NMDG solution before being transferred in a solution containing (in mM): 125 NaCl, 26 NaHCO_3_, 2 CaCl_2_, 2.5 KCl, 2 MgCl_2_, 0.8 NaH_2_PO_4_, and 10 D-glucose and equilibrated with 95% O_2_ - 5% CO_2_. Each slice was transferred to a submerged chamber mounted on an upright microscope (Olympus, BX51 WI) and neurons were visualized using differential interference contrast infrared video-microscopy.

### Electrophysiology

Whole-cell patch-clamp recordings were obtained from dLGN relay neurons. The external saline contained (in mM): 125 NaCl, 26 NaHCO_3_, 3 CaCl_2_, 2.5 KCl, 2 MgCl_2_, 0.8 NaH_2_PO_4_, and 10 D-glucose and equilibrated with 95% O_2_ - 5% CO_2_. Synaptic inhibition was blocked with 100 µM picrotoxin. In experiments where LTP-IE was induced with current pulses, excitatory synaptic transmission was also blocked with 2 mM kynurenate. Patch pipettes (5-10 MΩ) were pulled from borosilicate glass and filled with an intracellular solution containing (in mM) 120 K-gluconate, 20 KCl, 10 Hepes, 0.5 EGTA, 2 MgCl_2_, 2 Na_2_ATP and 0.3 NaGTP (pH 7.4). Recordings were performed with a MultiClamp-700B (Molecular Devices) at 30°C in a temperature-controlled recording chamber (Luigs & Neumann, Ratingen, Germany). The membrane potential was corrected for the liquid junction potential (−13 mV). In all recordings, access resistance was fully compensated using bridge balance and capacitance neutralization (>70%). dLGN neurons were recorded in current-clamp and input resistance was monitored throughout the duration of the experiments. Cells that display a variation >20% were discarded from final analysis. Voltage and current signals were low pass-filtered (10 kHz) and sequences of 2 s were acquired at 20 kHz with pClamp (Axon Instruments, Molecular Devices). Intrinsic excitability was tested with injection of current pulses (150-250 nA, 800 ms) to elicit 3-4 action potentials in control conditions at a frequency of 0.1 Hz. The amplitude of the current pulse was kept constant before and after induction of LTP-IE.

### Synaptic stimulation and induction of LTP-IE

Glass stimulating electrodes were filled with the extracellular medium. EPSPs were evoked in dLGN neurons with a pipette placed on the retino-thalamic endings of optic tract. Typically, EPSP amplitude reached its maximum value with stimulus intensity below 50-70 pA. The stimulus intensity was adjusted either to evoke an action potential by each EPSP (intensity: 50-100 µA) or on the opposite, no action potential (intensity: 20-40 µA). To mimic visual stimulation, trains of 15 synaptic stimulations at 40 Hz were applied during 10 minutes. In some cases, action potentials were evoked by short steps of current (2-5 ms, 1.0-2.5 nA) in the form of trains of 15 pulses at a frequency of 40 Hz during 10 minutes. The amplitude of the current pulse was chosen to elicit a single action potential by the current pulse (i.e., 15 action potentials per train). Each train occurred at a frequency of 0.1 Hz.

### Data analysis

Electrophysiological signals were analysed with ClampFit (Axon Instruments, Molecular Devices). Spikes were counted using Igor Pro software (Wavemetrics). Input resistance was calculated from voltage response to small negative current pulses (typically, −20 pA, 250 ms). LTP-IE was measured over a period of 10 minutes, 20 minutes after the beginning of the post-stimulation period.

### Drugs

1,2-Bis(2-aminophenoxy) ethane-*N*, *N*, *N*’, *N*’-tetraacetic acid (BAPTA) was added to the intracellular solution and was obtained from Tocris Bioscience. All other chemicals were bath applied. Picrotoxin and arachidonyl trifluoromethyl ketone (AACOCF3) were purchased from Abcam, DTx-I, DTx-k and 3,5-dichloro-N-[1-(2,2-dimethyl-tetrahydro-pyran-4-ylmethyl)-4-fluoro-piperidin-4-ylmethyl]-benzamide (TTA-P2) from Alomone and (3*R*, 4a*R*, 5*S*, 6*S*, 6a*S*, 10*S*, 10a*R*, 10b*S*)-5-(Acetyloxy)-3-ethenyldodecahydro-6,10,10b-trihydroxy-3,4a,7,7,10a-pentamethyl-1*H*-naphtho[2,1-*b*] pyran-1-one (forskolin), 9*R*,10*S*,12*S*)-2,3,9,10,11,12-Hexahydro-10-hydroxy-9-methyl-1-oxo-9,12-epoxy-1*H*-diindolo[1,2,3-*fg*:3’,2’,1’-*kl*] pyrrolo[3,4-*i*] [1,6] benzodiazocine-10-carboxylic acid, hexyl ester (KT5720) and nitrendipine were purchased from Tocris Bioscience.

### Immunostaining

Postnatal P11 and P25 wild-type Long Evans rats were deeply anesthetized and perfused with ice-cold 4% paraformaldehyde diluted in glucose 20 mM in phosphate buffer 0.1 M pH 7.4. Brains were removed from the skull and post-fixed overnight (o/n) at 4°C in the same fixative solution. 70 µm thick sections were cut using Leica VT1200S vibratome and then processed for immuno-histochemical detection. Brain slices were washed in DPBS (Gibco #14190-094) and incubated 30 min in sodium citrate 10 mM pH 8.5 at 80°C for antigen retrieval. Then there washed with phosphate buffer 0.1 M pH 7.4 (PB). Immunodetection was done in free-floating sections. Brain slices were treated with 50 mM NH_4_Cl for 30 min and incubated in blocking buffers to diminish nonspecific binding, using a first step PB with 1% (w/v) BSA (Sigma #A3059), 0.3% (v/v) TritonX100 (Merk #648463) for 30 min, then PB with 5% (w/v) BSA, 0.3% (v/v) Triton X-100 for 2 h. Slices were incubated overnight at 4°C with primary antibodies, mouse anti Kv1.1 clone K36/15 IgG2b (5 µg/ml) from Sigma-Aldrich and rabbit anti-ankyrinG (2.5 µg/ml), from Synaptic System (#386003), both diluted in incubation buffer (1% (w/v) BSA, 0.3% (v/v) Triton X-100 in PB). After extensive washing in incubation buffer, the secondary antibodies, Alexa 488 donkey anti-mouse (2 µg/ml; #715-546-151) and Alexa 594 donkey anti rabbit (2 µg/ml; #711-586-152) from Jackson Immunoresearch were incubated for 2 h at room temperature and then washed. After staining, the coverslips were mounted with Vectashield Vibrance (Vector H-1700-10).

### Confocal fluorescence microscopy and image analysis

Images were acquired on a confocal laser scanning microscope (LSM 780 Zeiss) using the same settings to compare intensities between experimental conditions P11 and P25 ages. All images were acquired using a Plan-Apo 63X /1.4 Oil-immersion objective lens. All confocal images were acquired at 0.38-μm z axis steps and with a 1,024 × 1,024-pixel resolution. Photons emitted from the two dyes were detected sequentially with one photomultiplier tube for each dye to minimize cross-talk. Images were prepared using Adobe Photoshop.

Analysis was performed using the ImageJ software (NIH). Images stacks were converted into single maximum intensity z axis projections. AIS was identified by AnkyrinG staining and was selected with threshold determination of fluorescent labeling area (red stain). After subtraction of the background, average gray value was measured on Kv1.1 staining within the selection (the sum of the gray values of all the pixels in the selection divided by the number of pixels). All statistical analyses were carried out in SigmaPlot.

### Statistics

Pooled data are presented as mean ± S.E.M. Statistical analysis was performed using Wilcoxon test or Mann-Whitney U test.

## Author contributions

DoD & MR conceived and supervised the project, MD, LFM, AW, EZ, CIB, & MR collected the data, MD, LFM, CIB, EZ, AA, SI, DaD, RB, BMP, MR & DoD analysed the data, DoD wrote the manuscript which was subsequently completed by all authors.

## Acknowledgments

Supported by INSERM, CNRS (to DoD), AMU (to MR), FRM (DVS20131228768 to DoD), ANR (LoGiK, ANR-17-CE16-0022 to DoD, ANR-21-CE16-0013 to DoD and RB), and NeuroSchool (to AW). This work received support from the French government under the France 2030 investment plan, as part of the Initiative d’Excellence d’Aix-Marseille Université – A*MIDEX, AMX-22-RE-AB-187 (to DoD) and AMX-22-RE-V2-007 (to DoD). We thank K Milton, O Toutendji & A Venture for excellent animal care.

---

**Figure S1.**
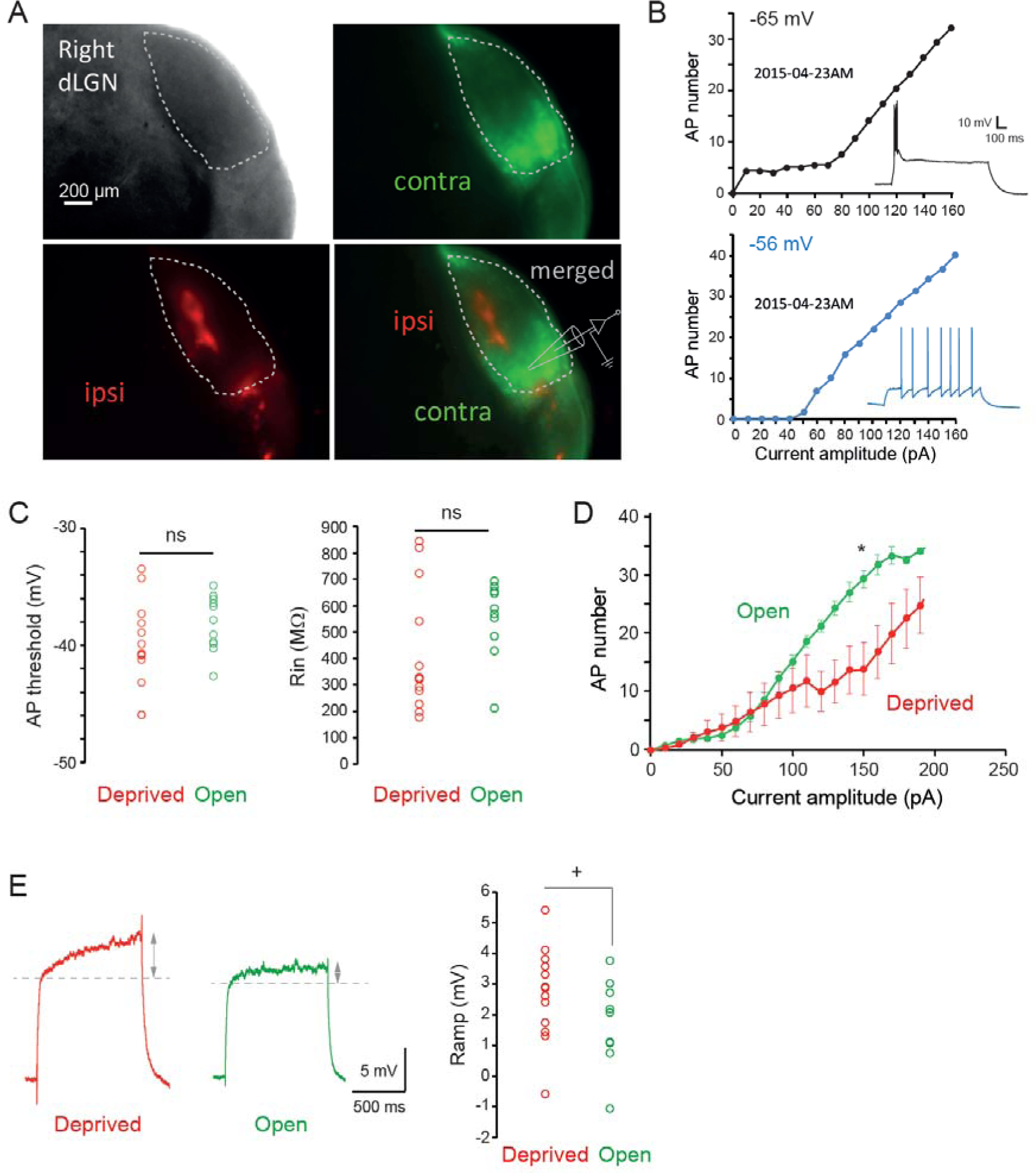
Definition of the retinal projections in the rat dLGN and changes in excitability. A. Labelling of the contralateral and ipsilateral retinal projections. B. Inactivation of the T-type calcium current reveals a delayed firing. At resting membrane potential (−65 mV, top), the rheobase is difficult to define because of the burst mediated by T-type calcium channels (see trace in response to 60 pA). However, when the same neuron is depolarized to −56 mV (bottom), the T-type current is inactivated (see trace in response to 60 pA) and the rheobase can be properly defined. C. Comparison of the input resistance (Rin, left) and of the AP threshold (right). D. Comparison of the input-output curves of open and deprived dLGN neurons measured at −65 mV. *, Mann-Whitney test, p <0.05. E. Comparison of subthreshold voltage ramps in neurons activated by the deprived and open eyes. The voltage ramp was found to be slightly larger in deprived neurons compared to open ones. Left, representative traces. Right, group data. Mann Whitney test, +, p<0.08.

**Figure S2.**
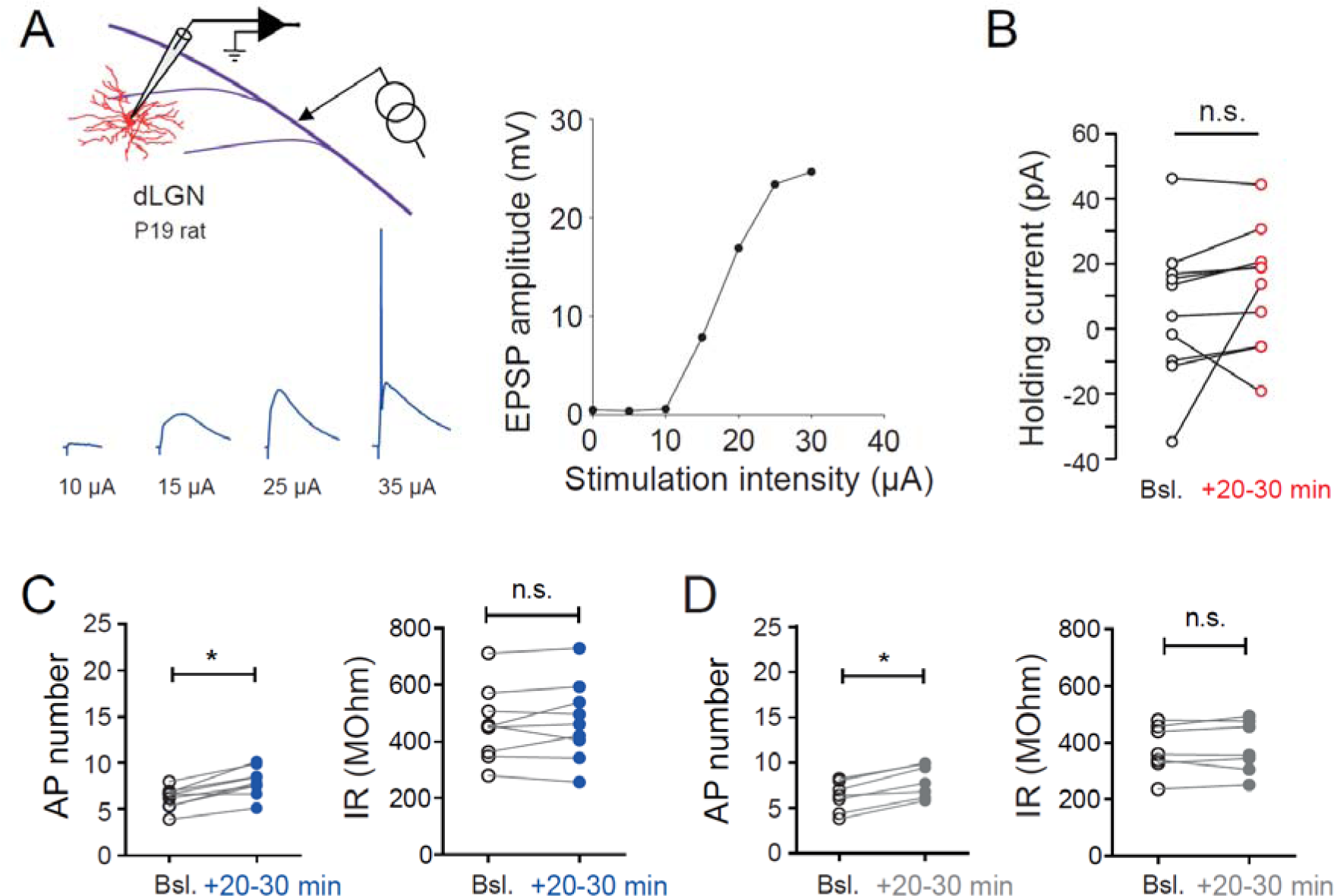
Synaptic responses, AP number and input resistance. A. Top left, recording and stimulation configuration. Bottom left, synaptic responses at different intensities. Note that at 35 µA, an AP is systematically evoked. Right, plot of the EPSP amplitude as a function of stimulus intensity. B. Holding current is not significantly changed. C & D. AP number and input resistance (IR) for subthreshold EPSPs (C) and no stimulation (D). Wilcoxon test, n.s., not significant, *, p<0.05.

**Figure S3.**
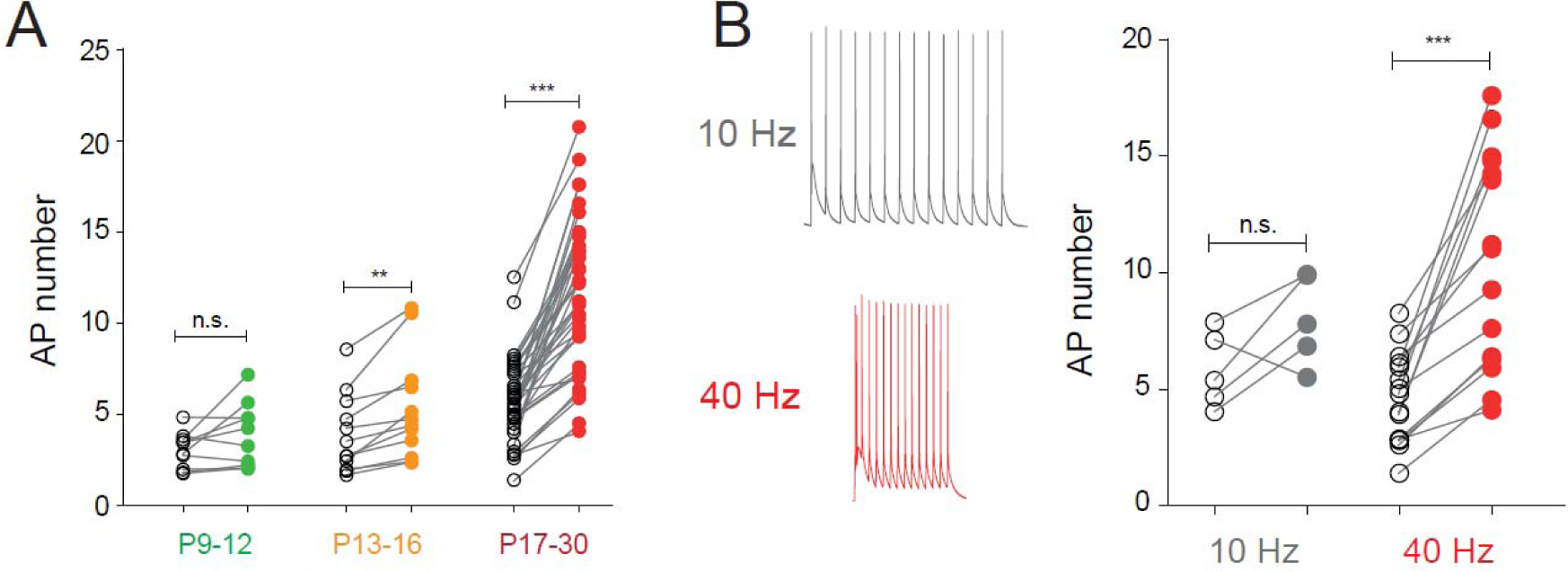
Age- and frequency-dependence of LTP-IE in dLGN neurons. A. Changes in spike number in each age category. Wilcoxon test, ns, not significant; **, p<0.01; ***, p<0.001. B. Frequency-dependence of LTPI-IE. Left, trace examples at 10 Hz (grey) and 40 Hz (red). Right, AP changes. Wilcoxon, ns, not significant; ***, p<0.001.

**Figure S4.**
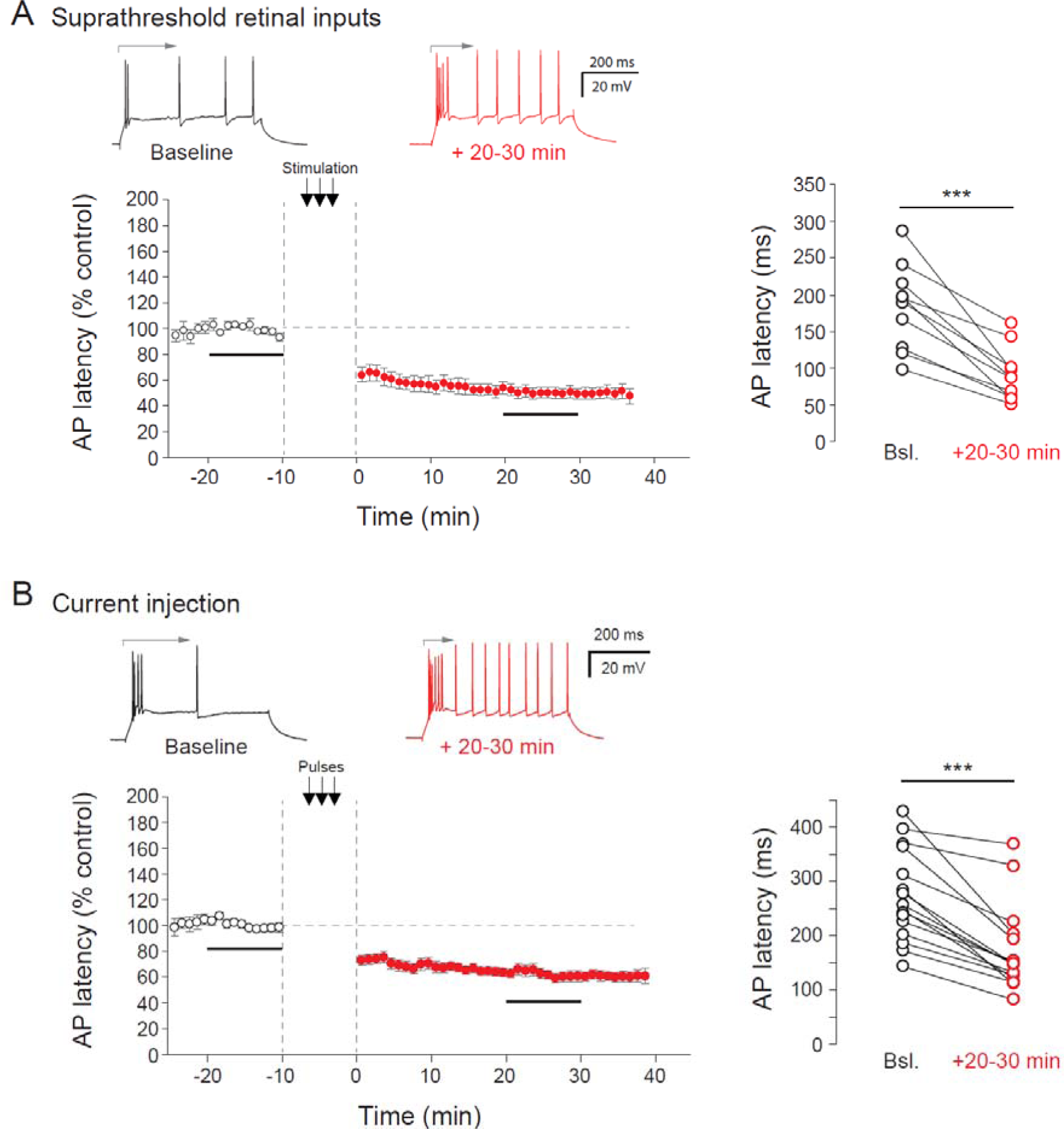
Reduction of the AP latency following induction of LTP-IE. A. Reduction of the AP latency of the first spike after the burst (indicated by an arrow) after induction of LTP-IE by stimulation of suprathreshold retinal inputs at 40 Hz. Left, time course. Right, group data. Wilcoxon test, ***, p<0.001. B. Reduction of the AP latency of the first spike after the burst (indicated by an arrow) after induction of LTP-IE by current pulse injection at 40 Hz. Left, time course. Right, group data. Wilcoxon test, ***, p<0.001.

**Figure S5.**
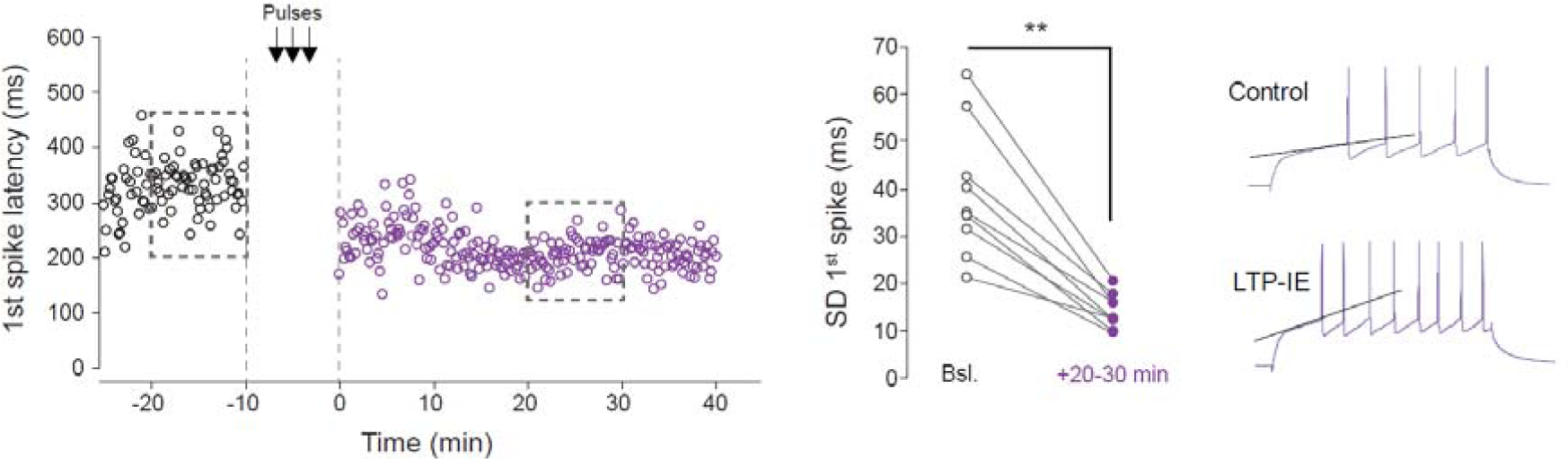
Reduced spike jitter following LTP-IE induction. Left, time-course of the first spike jitter in a neuron recorded in the presence of TTA-P2. Middle, group data. **, Wilcoxon test, p<0.01. SD, standard deviation. Right, slope changes before and after induction of LTP-IE.

**Figure S6.**
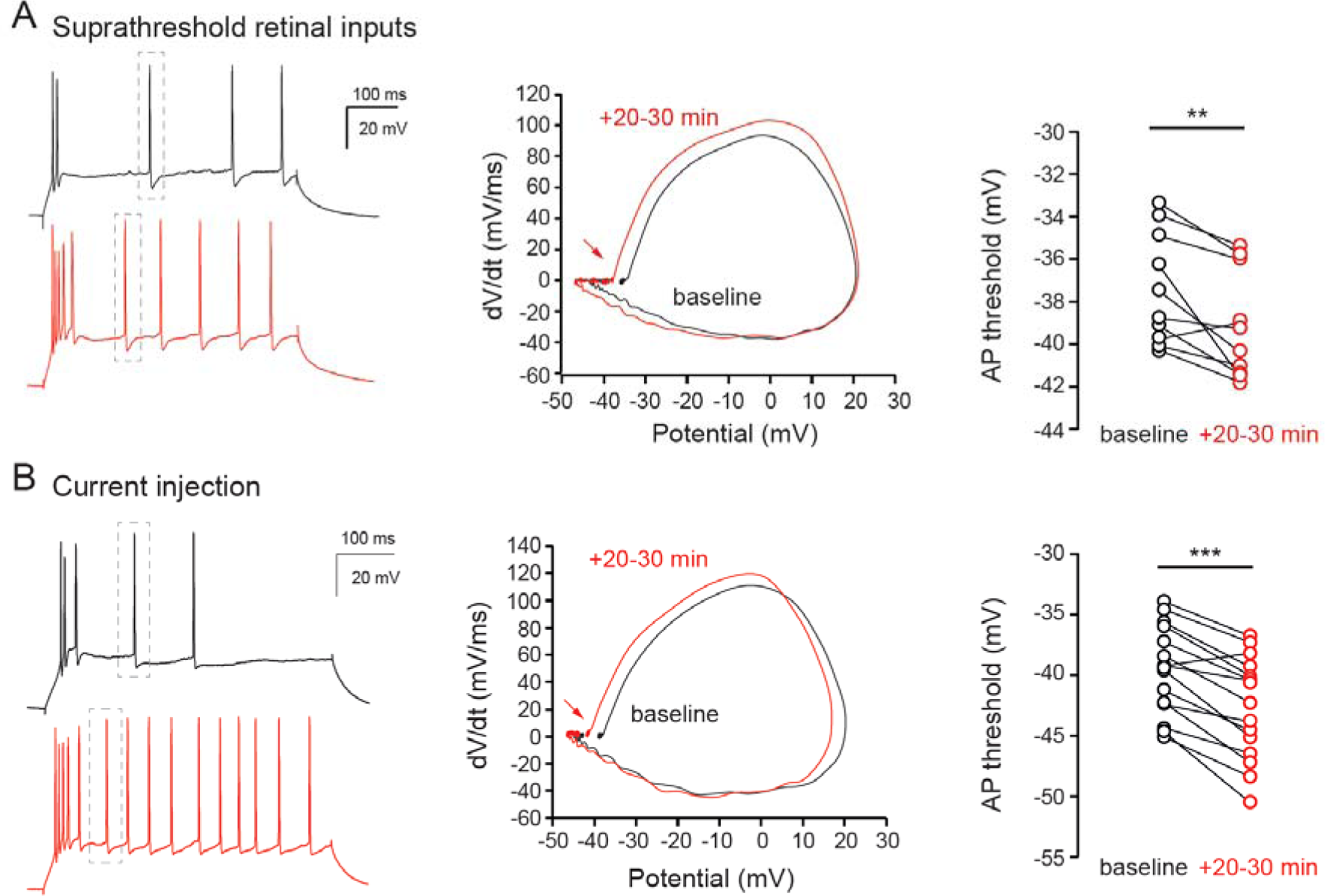
Hyperpolarization of the AP threshold after induction of LTP-IE in dLGN neurons. A & B. Hyperpolarization of the AP threshold after induction of LTP-IE with stimulation of suprathreshold retinal inputs (A) and by current injection (B). Left, representative traces. Middle, phase plots. Right, group data. Wilcoxon test, ** p<0.01; ***, p<0.001.

**Figure S7.**
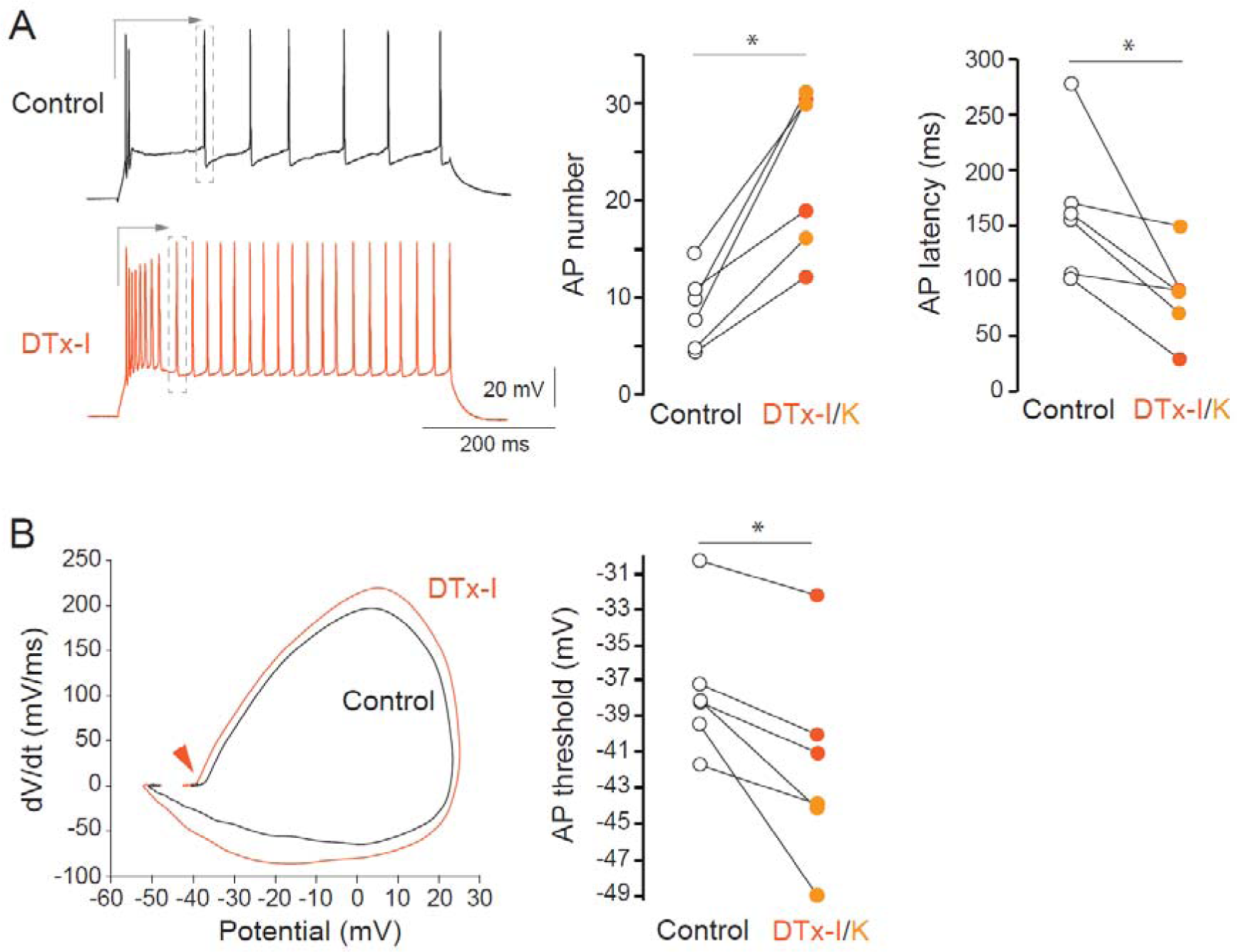
Action of DTx on firing properties of dLGN neurons. A. DTx increases evoked firing and reduces the delay of the first spike after the burst. Left, traces in control (black) and in DTx-I (orange). Right, pooled data for AP number and AP latency. Orange data points correspond to DTx-I and light-orange ones to DTx-K. B. DTx hyperpolarizes the AP threshold. Left, phase plots in control and DTx-I. Right, pooled data. Orange data points correspond to DTx-I and light-orange ones to DTx-K. Wilcoxon test, * p<0.05.

**Figure S8.**
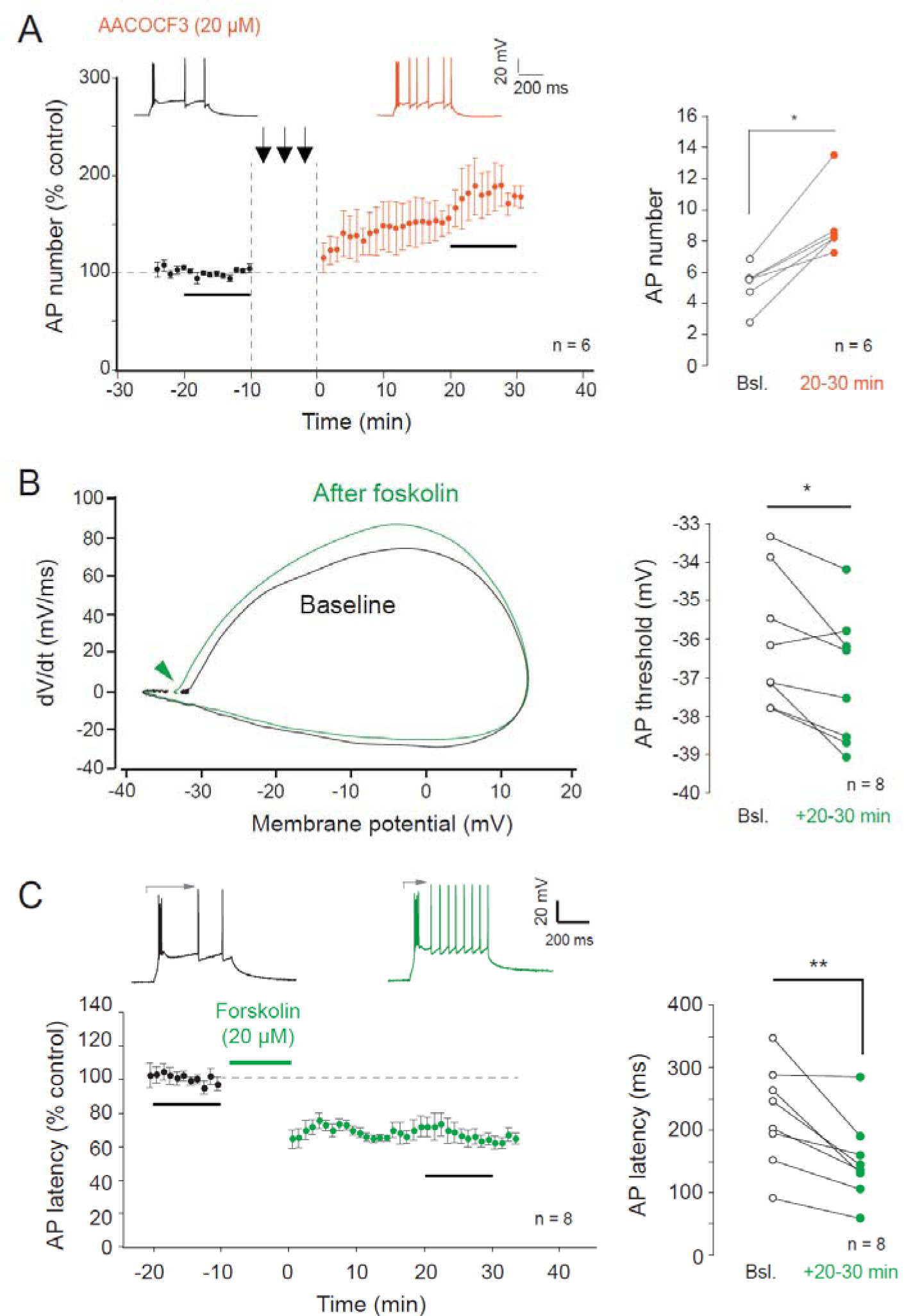
Induction of LTP-IE in the presence of an inhibitor of AA production and changes in spike threshold and latency following application of forskolin. A. Induction of LTP-IE in dLGN neurons is not blocked by the inhibitor of arachidonic acid synthesis, AACOCF3. Left, time course. Right, pooled data. Wilcoxon test, *, p<0.05. B. AP threshold hyperpolarization following forskolin application. Left, phase plots. Right, pooled data. Wilcoxon, *, p<0.05. C. Reduction of spike latency following application of forskolin. Left, time course. Right, group data. Wilcoxon test, **, p<0.01.

